# Genome assembly with variable-order de Bruijn graphs^⋆^

**DOI:** 10.1101/2022.09.06.506758

**Authors:** Diego Díaz-Domínguez, Pierfrancesco Martinello, Taku Onodera, Simon J. Puglisi, Leena Salmela

## Abstract

Choosing an order for constructing a *de Bruijn graph* (DBG) is a crucial step in *de novo* assembly, as no single value allows complete genome reconstruction. The *variable-order de Bruijn graph* (voDBG) addresses this limitation by combining DBGs of multiple orders in a single structure connected by contextual relationships. This representation enables new connections to be identified or ambiguities to be resolved during assembly. However, voDBGs currently lack a formal definition of contigs.

In this paper, we give the first formal definition of contigs for voDBGs. We show that, for a frequency range [*ℓ, h*] with *ℓ > h/*2, nodes whose labels occur with frequency *f* ∈ [*ℓ, h*] in the reads spell sequences of the genome with high probability under uniform sampling assumptions. We call these sequences (*ℓ, h*)-tigs. We also present an efficient algorithm to enumerate (*ℓ, h*)-tigs from a voDBG that accounts for homopolymer errors. Experiments on PacBio HiFi data show that our method significantly improves contiguity compared to fixed-order DBGs while remaining considerably lighter than full genome assemblers.

## 1 Introduction

Short-read assemblers have historically relied on *de Bruijn graphs* (DBGs) [15,27] to reduce computational costs. These methods decompose reads into *k*-mers, construct the corresponding DBG, and reconstruct the genome by traversing maximal unary paths. Although effective, their performance depends critically on the choice of *k*. Selecting a value that maximizes assembly quality is non-trivial [9]: small *k* values produce tangled graphs, whereas large *k* values lead to fragmentation due to variations in genome complexity and sequencing coverage.

Advances in DNA sequencing have produced longer and more accurate reads. PacBio HiFi reads, for example, are typically 10-20 kbp long with error rates below 1% [31]. These improvements, together with fast heuristics for approximate alignment [3,25,19], have facilitated the use of overlap–layout–consensus (OLC) strategies [21] in genome assembly. OLC methods often yield more accurate reconstructions than DBG-based approaches but require the costly computation of suffix-prefix overlaps between reads, a task further complicated by homopolymer-length errors. HiFi assemblers based on this paradigm include Peregrine [10], HiCanu [23], and hifiasm [6]. Despite alignment heuristics, constructing OLC graphs from HiFi reads remains expensive, motivating renewed interest in DBG-based approaches that scale more gracefully.

With HiFi reads, a DBG with a large *k* can disentangle the graph without substantially fragmenting it and thus improve contiguity. The challenge lies in the efficient construction and representation of such large-*k* DBGs. Some methods progressively increase *k*, as in IDBA [26] and SPAdes [2], whereas others selectively increase *k* in complex regions, such as LJA [1]. Other strategies reduce graph size through sparsification or compression: MBG [29] constructs a sparse DBG from a subset of *k*-mers, while mdBG [13] performs assembly in minimizer space. More generally, models supporting variable-length contexts have been proposed. Manifold DBGs [20] allow arbitrary substrings as node labels, and variable-order DBGs (voDBGs) [4] represent DBGs of all orders up to *K*.

Among these alternatives, voDBGs provide an appealing framework for large-*k* assembly. They retain information from all orders up to *K* and are closely related to compressed indexes with suffix-tree functionality [22]. Such structures admit efficient construction [12,24] and compact representation [5], suggesting that voDBGs could be built and stored efficiently.

However, a fundamental theoretical gap remains. In fixed-order DBGs, contigs are defined by node in- and out-degrees. In a voDBG, edges encode both sequence extensions and changes in context length, so purely topology-based notions of contigs, such as unitigs and omnitigs [30], do not apply directly. No formal definition of contigs has yet been established for voDBGs, and no assembly framework has yet been developed for this model.

### Our contribution

We present the first definition of contigs for a voDBG. We consider a frequency-restricted subgraph of the voDBG and show that, when node frequencies are limited to an interval [*ℓ, h*] with *ℓ > h/*2, the subgraph corresponds to a set of strings that we call (*ℓ, h*)-tigs. We analyze how the choice of *ℓ* and *h* affects the contiguity and accuracy of the assembly and provide a method for choosing appropriate values of *ℓ* and *h*. We further exploit connections between the voDBG and compressed indexes to develop an efficient, homopolymer-aware algorithm for enumerating (*ℓ, h*)-tigs. We implement our ideas in a tool called `Ryu` and show that our approach achieves better contiguity relative to fixed-order DBG assemblers. While full-featured assemblers still achieve higher contiguity by handling residual assembly inconsistencies, `Ryu` shows comparable accuracy with lower computational costs.

## 2 Preliminaries

### DNA strings

A *read R*[1..*n*] is a sequence over the alphabet Σ = {`a`, `c`, `g`, `t`}. Standard notions of suffix, prefix, and substring apply. A *run* is a maximal substring of identical symbols. A run of length greater than one is called a *homopolymer*. The run-length encoding of 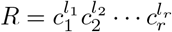 is *rle*(*R*) = (*c*_1_, *l*_1_), (*c*_2_, *l*_2_), …, (*c*_*r*_, *l*_*r*_).

The *complement c* : Σ → Σ maps `a` ↔ `t` and `c` ↔ `g`; the *reverse complement* of *R* is denoted 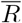. A *read collection* is a multiset ℛ= *{R*_1_, *R*_2_, …, *R*_*q*_*}* of DNA strings. Its extended version 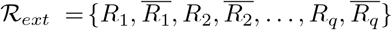 contains each read together with its reverse complement.

### de Bruijn graphs

The node-centric *de Bruijn graph* (DBG) of order *k* [11] is a labeled graph *G* = (*V, E*) in which each node *u* ∈ *V* represents a distinct *k*-mer *label*(*u*) ∈ Σ^*k*^. There is an edge (*u, v*) ∈ *E* iff *label*(*u*)[2..*k*] = *label*(*v*)[1..*k* − 1]. The edge (*u, v*) is labeled with *label*(*u*)[*k*].

Given a read collection ℛ, the induced graph *G*^ℛ^ ⊆ *G* contains only those nodes whose labels occur in ℛ and those edges (*u, v*) such that *label*(*u*)·*label*(*v*)[*k*] occurs in ℛ. We define the operator *f* (*u*) to denote the frequency of *S* = *label*(*u*) in ℛ. We use *f* (*u*) and *f* (*S*) interchangeably.

### Variable-order DBGs

The *variable-order de Bruijn graph* (voDBG) [4] of ℛ combines the DBGs of all orders 1 ≤ *k* ≤ *ρ*, where *ρ* is the maximum read length. Its node set consists of all distinct substrings of ℛ, grouped by length.

For a node *u* of order *k*:

*Right-extension edge:* (*u, v*) to a node *v* of order *k* + 1 exists if *label*(*u*) = *label*(*v*)[1..*k*]. The edge is labeled with *label*(*v*)[*k* + 1].

*Left-contraction edge:* (*u, w*) to a node *w* of order *k* − 1 exists if *label*(*u*)[2..*k*] = *label*(*w*).

Left extension and right contraction edges are defined symmetrically.

## 3 Our framework

Our assembly method traverses the voDBG of the reads along paths that alternate extension and contraction edges. Extensions refine sequence context and continue while possible; contractions restore context when extensions stop. A contig is formed by the label of the starting node followed by the labels of the visited extension edges.

Extensions decrease frequency and contractions increase it, so we restrict the traversal to nodes whose frequencies lie within the sequencing-coverage interval.

### 3.1 Description of contigs in the voDBG

In this section, we describe our formal concept of contigs in a voDBG *G* = (*V, E*) induced from an *ideal* read collection ℛ. Ideal means that the reads were sampled uniformly at random from one strand of a haploid genome *W* with coverage [*ℓ, h*], are error free, and have uniformly distributed endpoints across *W*. We also assume the repeats of *W* are fully resolved within the reads.

Although idealized, this model captures the dominant behavior of HiFi data. Section 3.2 discusses how to deal with more realistic scenarios.

#### Definition 1 (Frequency-restricted voDBG)

*Given an interval* [*ℓ, h*], *a frequency-restricted voDBG G*_*ℓ,h*_ ⊆ *G retains each node u such that ℓ* ≤ *f* (*u*) ≤ *h and each edge* (*u, v*) *such that ℓ* ≤ *f* (*u*) ≤ *h and ℓ* ≤ *f* (*v*) ≤ *h*.

#### Lemma 1.

*When h/*2 *< ℓ < h, each node in G*_*ℓ,h*_ *has at most one outgoing and at most one incoming extension edge. Similarly, each node has at most one outgoing and at most one incoming contraction edge*.

*Proof*. Suppose a node *v* ∈ *V* of order *k* + 1 has two incoming extension edges (*u*_1_, *v*) and (*u*_2_, *v*). Both *u*_1_ and *u*_2_ are the prefixes *label*(*v*)[1..*k*] = *label*(*u*_1_) = *label*(*u*_2_) and therefore *u*_1_ = *u*_2_. Thus, *v* has one incoming extension edge.

Now suppose a node *u* ∈ *V* of order *k* has two outgoing extension edges (*u, v*_1_) and (*u, v*_2_) with *label*(*v*_1_)[*k* + 1] = *label*(*v*_2_)[*k* + 1]. Since *label*(*v*_1_)[1..*k*] ≠ *label*(*v*_2_)[1..*k*] = *label*(*u*), it follows that *f* (*u*) ≥ *f* (*v*_1_) + *f* (*v*_2_).

Because *v*_1_, *v*_2_ ∈ *V*, we have *f* (*v*_1_), *f* (*v*_2_) ≥ *ℓ*, and therefore *f* (*u*) ≥ 2*ℓ*. Since *ℓ > h/*2, we also obtain 2*ℓ > h*, and hence *f* (*u*) ≥ 2*ℓ > h*. This inequality contradicts *u* ∈ *V*, which requires *f* (*u*) ≤ *h*. Therefore, *u* has at most one outgoing extension edge.

The statement for contraction edges follows by symmetry.

Notice the condition *ℓ > h/*2 is tight: if *ℓ* ≤ *h/*2, branching nodes may exist. From now on, we will assume *ℓ > h/*2 holds in *G*_*ℓ,h*_.

#### Lemma 2.

*Extension and contraction edges in G*_*ℓ,h*_ *commute: (1) if* (*u, v*) *is an extension edge labeled c and* (*u, w*) *is a contraction edge, there exists a node x such that* (*v, x*) *is a contraction edge and* (*w, x*) *is an extension edge labeled c and (2) if* (*w, x*) *is an extension edge labeled c and* (*v, x*) *is a contraction edge, there exists a node u such that* (*u, w*) *is a contraction edge and* (*u, v*) *is an extension edge labeled c. See Figure 1*.

*Proof*. (1) Let *w, u* and *v* be nodes with orders *k* − 1, *k* and *k* + 1, respectively, such that the extension (*u, v*) and the contraction (*u, w*) exist. Then it follows *label*(*v*) = *label*(*u*) · *c* and *label*(*w*) = *label*(*u*)[2..*k*].

Define a node *x* with *label*(*x*) = *label*(*w*) ·*c* = *label*(*u*)[2..*k*] ·*c*. This implies that (*w, x*) is an extension edge labeled *c*. Furthermore, since *label*(*v*) = *label*(*u*) ·*c*, a left contraction yields *label*(*u*)[2..*k*] ·*c* = *label*(*x*), so (*v, x*) is a contraction edge.

An extension decreases frequency and a contraction increases it, so we have *f* (*v*) ≤ *f* (*x*) ≤ *f* (*w*). Because *u, v, w* ∈ *V* and their frequencies lie in [*ℓ, h*], it follows that *f* (*x*) ∈ [*ℓ, h*], hence *x* ∈ *V*. (2) follows by symmetry.

**Fig. 1.**
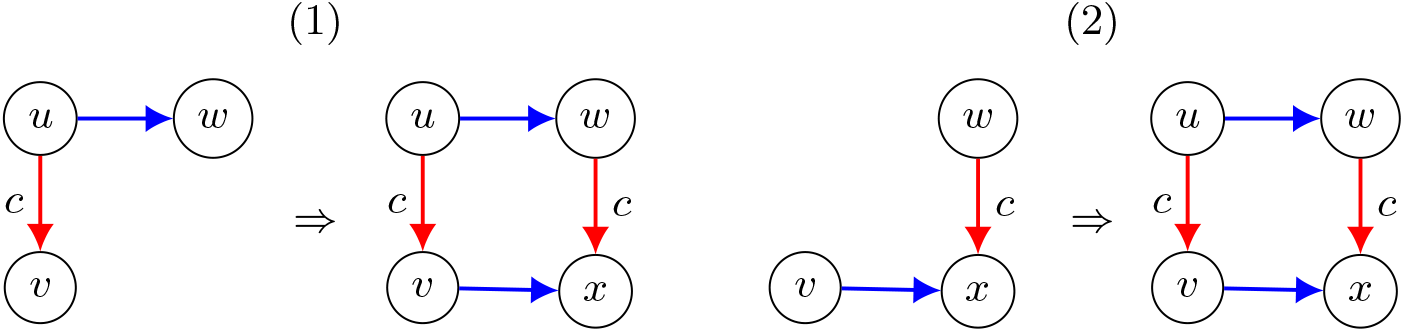
Lemma 2. Red edges are extensions and blue edges are contractions.

Because contraction edges are non-branching (Lemma 1), they partition *G*_*ℓ,h*_ into maximal directed paths, which we call *contraction paths*.

#### Lemma 3.

*Let U* = (*u*_*i*_)_1≤*i*≤*n*_ *and P* = (*p*_*i*_)_1≤*i*≤*m*_ *be distinct contraction paths in G*_*ℓ,h*_. *If there is an extension edge* (*u*_*i*_, *p*_1_) *with* 1 ≤ *i* ≤ *n labeled with c, then for every* 0 ≤ *j* ≤ *n* − *i, there is also an extension edge* (*u*_*i*+*j*_, *p*_1+*j*_) *labeled c*.

*Proof*. Follows from successive applications of Lemma 2.

Along a contraction edge (*u*_*i*_, *u*_*i*+1_), if every occurrence of *label*(*u*_*i*+1_) in ℛ occurs within *label*(*u*_*i*_), then *f* (*u*_*i*_) = *f* (*u*_*i*+1_). In this case, *u*_*i*_ and *u*_*i*+1_ have the same right extension, so the extension edge is unchanged. This idea is illustrated in Figure 2 with dots at the ends of *U* and *P*.

**Fig. 2.**
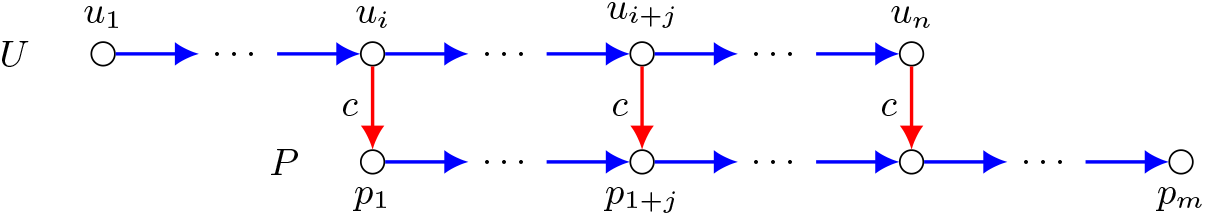
Lemma 3

Lemma 3 allows us to lift *G*_*ℓ,h*_ to a meta-graph:

#### Definition 2 (voDBG meta-graph)

*The meta-graph* 𝒢= (𝒱, ℰ) *of G*_*ℓ,h*_ *is defined as follows:*

*Each maximal contraction path U in G*_*ℓ,h*_ *corresponds to a node in* 𝒱. *Since contraction strictly decreases the order, U contains a unique node of maximal order. The sequence label*(*U*) *is defined as the label of that node*.

*Two meta-graph nodes U, P* ∈ 𝒱 *are connected by an edge* (*U, P*) ∈ ℰ *labeled c iff there exists an extension edge labeled c in G*_*ℓ,h*_ *between U and P*.

By Lemmas 1 and 3, each node in 𝒢 has indegree and outdegree at most one. Consequently, connected components of 𝒢 are directed paths or cycles. We use this structure to define (*ℓ, h*)-tigs analogously to unitigs in fix-order DBGs:

#### Definition 3 ((*ℓ, h*)-tigs)

*Let* 𝒫= (*U*_1_, *U*_2_, …, *U*_*q*_) *be the nodes of a connected component of* 𝒢, *ordered so that* (*U*_*i*_, *U*_*i*+1_) ∈ ℰ *for i* ∈ [1, *q* − 1]. *If* 𝒫 *is a directed cycle, the choice of U*_1_ *is arbitrary and* (*U*_*q*_, *U*_1_) ∈ ℰ. *If* 𝒫 *is a directed path, U*_1_ *has indegree* 0 *and U*_*q*_ *has outdegree* 0. *An* (*ℓ, h*)*-tig is the string formed by label*(*U*_1_) *followed by the labels of the edges* (*U*_*i*_, *U*_*i*+1_) *in* 𝒫.

#### Spelling (*ℓ, h*)-tigs

We spell (*ℓ, h*)-tigs by traversing 𝒢 = (𝒱, ℰ). For each *U* ∈ 𝒱with indegree 0, we initialize *L* = *label*(*U*) and follow successive extension edges (*U*_*i*_, *U*_*i*+1_) ∈ ℰ, appending their labels to *L*, until reaching a node with outdegree 0. The resulting *L* is an (*ℓ, h*)-tig. For components that are directed cycles, we start from an arbitrary node and traverse the cycle once. After visiting all connected components of ℰ, the algorithm outputs one string per component.

#### Connection with genome assembly

Under the ideal conditions:

1. For each *U* ∈ 𝒱, *label*(*U*) is unique in *W*. Otherwise *f* (*label*(*U*)) *> h*, contradicting the frequency restriction.
2. If (*U, P*) is labeled *c* and *S* = *label*(*U*) · *c*, then *f* (*S*) *< f* (*label*(*U*)) reflects reads ending at *label*(*U*) and cannot extend by *c*.
3. Let *u*_*i*_ ∈ *U* be the leftmost node exposing the extension toward *P*. Then *f* (*label*(*u*_*i*_)) *> f* (*label*(*U*)) reflects additional reads whose longer prefix enables the extension.

##### Lemma 4 (Genome reconstruction).

*Under ideal conditions of* ℛ, *the set of* (*ℓ, h*)*-tigs represents a reconstruction of the underlying genome W*.

*Proof*. Extension edges correspond to substrings observed in the reads, while contraction edges represent shorter contexts shared among reads. The restriction *ℓ > h/*2 (Lemma 1) removes competing contexts in *G*_*ℓ,h*_, so these shared contexts correspond, under ideal conditions, to overlaps that allow extensions beyond single reads. The meta-graph ℛ reflects this property: each node has at most one incoming and one outgoing extension edge. Consequently, each component of ℛ spells a maximal sequence occurring in *W*. If the component is a cycle, the genomic region is a circular chromosome. Hence the (*ℓ, h*)-tigs reconstruct *W*.

#### Deviations from the ideal model

In practice, reads are affected by sequencing errors, unsolved repeats, non-uniform coverage, and (in polyploid genomes) allele-specific coverage variation. These factors cause (*ℓ, h*)-tigs to be interrupted (fragmentation) or deviate from true genomic substrings (misassembly).

*Fragmentation* occurs when the true genomic continuation of a (*ℓ, h*)-tig is removed from *G*_*ℓ,h*_ because its coverage drops below *ℓ*.

*Misassemblies* arise when the frequency of case (3) varies so that the number of reads supporting the correct extension decreases and matching reads from other contexts keep the frequency within [*ℓ, h*]. This problem combines a lack of coverage with spurious overlaps generated by sequencing errors and/or repeats. Figure 3 shows this scenario.

**Fig. 3.**
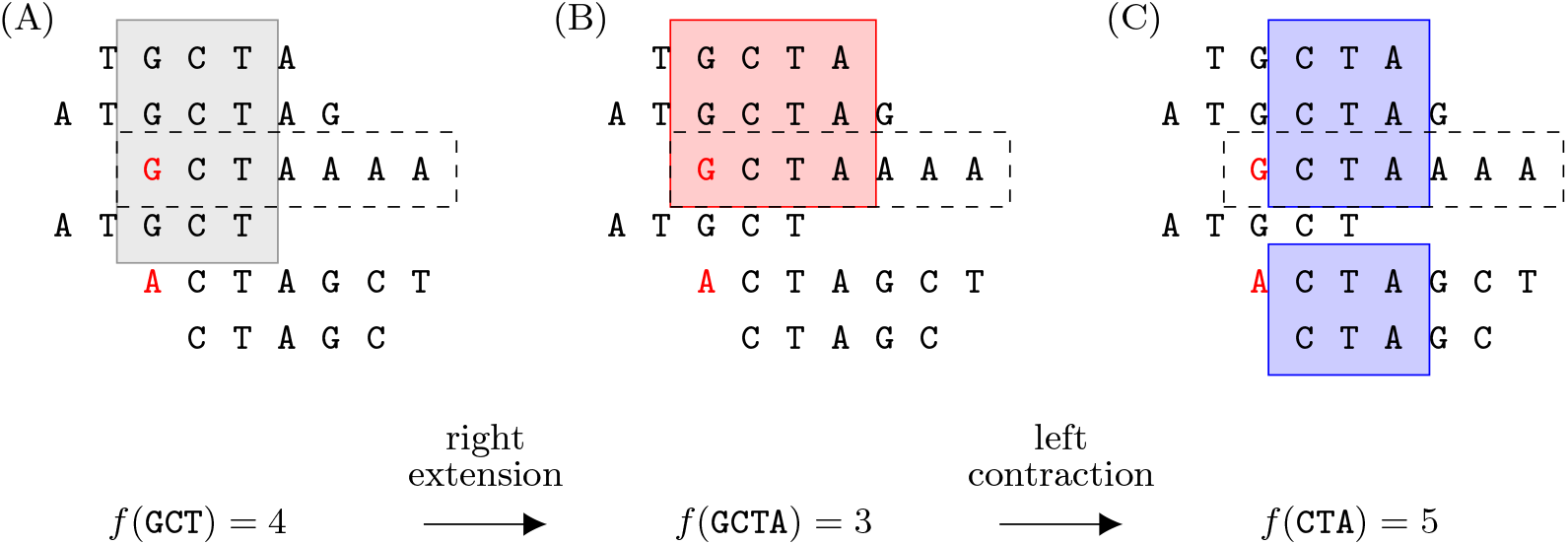
Genome assembly as a sequence of extensions and contractions. Red symbols indicate mismatches. The dashed read comes from another genomic region and creates a spurious match. In (A), `GCT` loses one occurrence due to a mismatch in `ACTAGCT`, and gains an occurrence from the spurious overlap. Extending `CTA` in (C) with `AAA` would result in a misassembly.

The choice of [*ℓ, h*] therefore balances these effects: larger values of *ℓ* remove spurious connections but increase fragmentation, while smaller values increase the connectivity at the cost of higher misassembly risk.

### 3.2 Determining *ℓ* and *h*

When the reads in ℛ do not satisfy the ideal conditions of Section 3.1, the interval [*ℓ, h*] must balance misassembly and fragmentation.

#### Modeling fragmentation

Let *δ* = *h* − *ℓ*+1 denote the maximum tolerated frequency fluctuation. Connectivity is disrupted when a contraction or extension changes frequency by at least *δ*.

Consider a substring *S* = *W* [*i*..*i*+*k*−1]. If *δ* reads start at *i*, then contraction from *cS* = *W* [*i*−1..*i*+*k*−1] yields *f* (*S*) ≥ *f* (*cS*) + *δ > h*, deleting the edge from *cS* to *S*. Symmetrically, if *δ* reads end at *i* + *k* − 1, the extension *Sc* = *W* [*i*..*i* + *k*] yields *f* (*Sc*) ≤ *f* (*S*) − *δ < ℓ*, deleting the edge from *S* to *Sc*.

Sequencing errors transfer occurrences of *S* in ℛ to other strings *S*^*′*^, decreasing *f* (*S*) and increasing *f* (*S*^*′*^). Assuming error rate *e*, coverage *C*, and genome size *w*, the expected number of such transfers is *eCw*.

The problem of choosing [*ℓ, h*] therefore reduces to a balls-into-bins model with |ℛ|+ *eCw* balls (effective endpoints) and *w* bins (genomic positions). By Raab and Steger [28], if

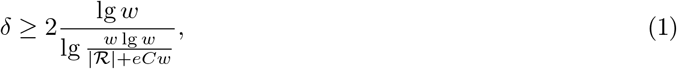

then with high probability no position of *W* is the start or end of *> δ* reads.

#### Modelling misassemblies

Suppose *S* has two genomic extensions *Sc*_1_ and *Sc*_2_. Under uniform coverage their read counts follow a binomial distribution with parameter 0.5. A misassembly occurs if one extension has frequency *i* ∈ [*ℓ, h*] (kept) while the other *f* (*S*) − *i < ℓ* (discarded). The probability of this happening is 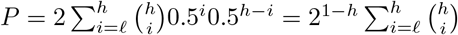.

Let *d* = *ℓ − h/*2 denote the deviation from the mean. By Chernoff bound, 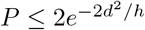. If a misassembly probability at most *ϵ* is tolerated, then

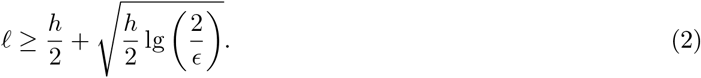

This analysis applies to the two-extension case. Combining Equations 1 and 2, we require

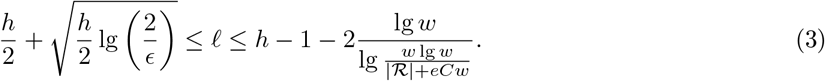

We choose the smallest *h* for which an integer *ℓ* satisfies Equation 3, ensuring that (*ℓ, h*)-tigs are neither fragmented nor prone to misassembly.

## 4 Practical genome assembler

We present a practical long-read assembler based on (*ℓ, h*)-tigs. Because sequencing samples both DNA strands, the input is the extended collection ℛ_*ext*_ containing each read and its reverse complement. We assume suitable values of [*ℓ, h*] (Section 3.2) computed from ℛ and doubled for ℛ_*ext*_.

Long-read technologies often misestimate homopolymer lengths, producing spurious overlaps and potential misassemblies. We incorporate a simple heuristic to mitigate this issue.

### Preparing the data

Each read *R* ∈ ℛ_*ext*_ is run-length encoded as *rle*(*R*) = (*c*_1_, *l*_1_)(*c*_2_, *l*_2_), …, (*c*_*r*_, *l*_*r*_). We separate it into its *symbol sequence c*_1_, *c*_2_, …, *c*_*r*_ and *length sequence l*_1_, *l*_2_, …, *l*_*r*_. Let ℛ_*c*_ be the collection of symbol sequences.

We index ℛ_*c*_ using a compressed structure *I* with suffix-tree functionality (see Section 5) that supports navigational queries in the voDBG *G*_*ℓ,h*_ = (*V, E*) of ℛ_*c*_. The length sequences are integrated into *I* so that for each node *u* ∈ *V* we can query

- *hp*(*u*): return the list of length sequences corresponding to the uncompressed occurrences of *label*(*u*) in ℛ_*ext*_.

The index *I* maps all occurrences of a substring *S* in ℛ_*ext*_ to the same node *u* in *G*_*ℓ,h*_ based on their symbol sequence, ignoring homopolymer length differences. This removes spurious overlaps caused by homopolymer errors. However, different strings *S* ≠ *S*^*′*^ with the same symbol sequence may also collapse to the same node. The lists returned by *hp*(*u*) allow us to detect such cases: homopolymer errors produce small deviations around the true lengths, whereas multiple strings typically generate distinct length distributions.

Because reverse complements are included, the meta-graph 𝒢= (𝒱, ℰ) contains two components per chromosome, one for each DNA strand. For a node *U* ∈ 𝒱 we denote by *Ū* the node labeled with the reverse complement of *label*(*U*). We also define *hp*(*U*) = *hp*(*u*_1_), where *u*_1_ is the highest-order node of *U*.

### Assembly

The algorithm selects a pair of nodes *U, Ū* ∈ 𝒱 with indegree and outdegree 0, respectively. These correspond to the starting and ending points of complementary components of 𝒢.

It reconstructs an uncompressed sequence for *S* = *label*(*U*) using the list ℒ= *hp*(*U*). For each position *i* in *S*, the homopolymer length is estimated as the median of the lengths at position *i* in ℒ. If the standard deviation exceeds a threshold *τ*, the length at position *i* is marked as *fuzzy*.

After processing all positions, if the fraction of fuzzy lengths exceeds a threshold *µ*, (*U, Ū*) is discarded and the algorithm moves to another component of 𝒢. Otherwise, the algorithm spells the (*ℓ, h*)-tig starting at *U*. The uncompressed length of each extension label *c* is estimated in the same way using the median.

Once the (*ℓ, h*)-tig is completed, the nodes of *U* and *Ū* are marked as processed, and the algorithm continues with the next unvisited pair (*U, Ū*).

## 5 Experiments

### Implementation details

We implemented our (*ℓ, h*)-tig assembler (Section 4) in `C++` as `Ryu`^1^. The tool builds a compressed FMD-index [18] over the symbol sequences ℛ_*c*_. The index uses the bidirectional BWT [17], computed with `grlBWT` [12]. Traversing BWT ranges corresponding to suffix tree nodes enumerates voDBG nodes *u* and their frequencies. Run-length encoding lengths from ℛ_*ext*_ are stored in BWT order to support *hp*(*u*) queries, enabling homopolymer reconstruction. `Ryu` then visits the connected components of the metagraph 𝒢 via BWT backward search and spells (*ℓ, h*)-tigs as described in Section 4.

`Ryu` currently supports limited parallelization, using up to four threads. To distribute work among them, the FMD-index is divided into ranges, which are then assigned to threads to compute the (*ℓ, h*)-tigs in parallel. However, without first processing the FMD-index, it is difficult to ensure that each thread receives a similar workload. Therefore, we adopt a simpler heuristic that partitions the data structure according to the DNA alphabet.

### Datasets

We used three PacBio HiFi read collections: *E. Coli* (ECOLI), *S. Cerevisiae* (YEAST), and the human cell line CHM13 (HUMAN), all being either complete or near haploid genomes. We assembled these datasets with different methods and evaluated the output using `Quast` [14]. Table 1 lists the SRA accessions of the reads and the reference genomes used for evaluation.

**Table 1.** Species and reference genomes used in the experiments. The reference genomes were downloaded from the NCBI database: https://www.ncbi.nlm.nih.gov/datasets/genome.

| Scientific name | Dataset | SRA accession | Ref. genome GenBank acc. | Genome size (Mbp) | Seq. coverage |
| --- | --- | --- | --- | --- | --- |
| *E. Coli* | ECOLI | SRR10971019 | GCA_000005845.2 | 4.64 | 300 |
| *S. Cerevisiae* | YEAST | SRR34579673 | GCA_051942865.1 | 11.36 | 676 |
| *H. Sapiens* | HUMAN | SRR11292121 | GCA_009914755.1 | 3,117.28 | 9.57 |

### Competitor tools

We compared `Ryu` with `Bcalm2`^2^ [8], `Flye`^3^ [16], and `Hifiasm`^4^ [7] for assembly. `Bcalm2` outputs unitigs from a fixed-order DBG, `Flye` constructs A-Bruijn and repeat graphs, and `Hifiasm` follows an OLC strategy based on minimizer-detected overlaps. `Bcalm2` and `Ryu` only spell sequences from the graph, whereas `Hifiasm` and `Flye` perform full genome assembly.

### Experimental setup

For `Ryu`, we varied *ℓ* (9 − 24) and *h* (16 − 32), and then calculated the theoretical optimal [*ℓ, h*] for each dataset (Section 3.2) with confidence rate 0.95. To increase assembly speed, we omitted (*ℓ, h*)-tigs with length smaller than 1000 base pairs using `-r 1000` (see discussion in Section 5.1). For `Bcalm2`, *k* ranged from 2^4^ to 2^8^, and the abundance threshold (`–abundance-min`) from 7 to 15. `Flye` and `Hifiasm` were run with default parameters. The experiments were executed with 24 threads to allow a comparison of memory usage and time between the tools. The only exception was `Ryu`, which used up to 4 threads due to implementation limitations.

Figure 4 illustrates the tradeoffs between assembly quality and the choice of (*ℓ, h*) (exact values are reported in Table A1. Assembly statistics for the compared tools are summarized in Table 2, while peak memory usage and runtime are listed in Table 3. For `Bcalm2`, we report the configuration with the fewest misassemblies in each dataset. For `Ryu`, we highlight two configurations: *best assembly* (`Ryu` ⋆), which achieves the highest N50 and genome coverage, and *best performance* (`Ryu` ▲), which completes the assembly of reads in the shortest time.

**Fig. 4.**
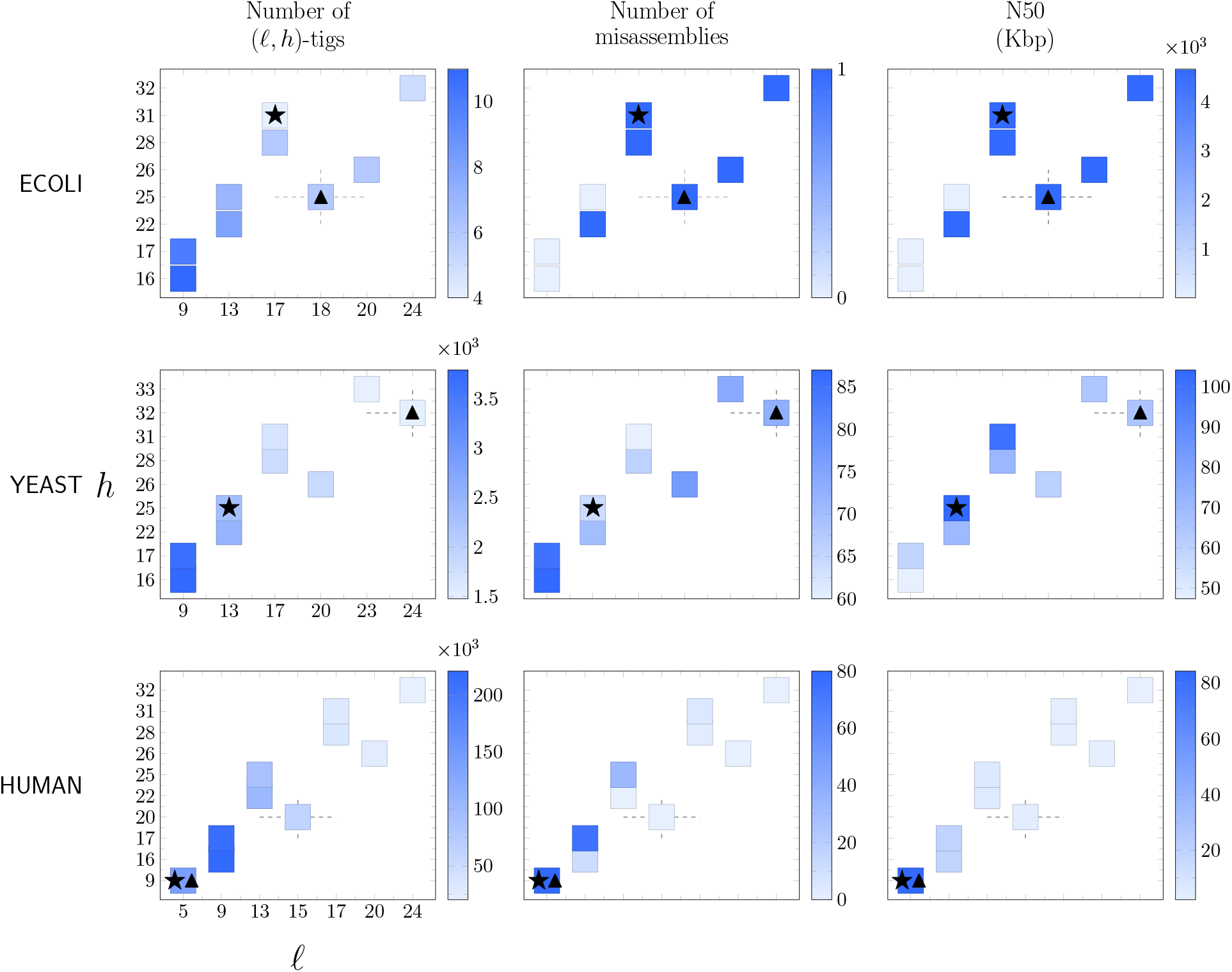
Effect of *ℓ* and *h* on the assembly. Each row corresponds to a dataset. The x-axis shows *ℓ* and the y-axis shows *h*. Axes are displayed only in the first column of each row because they are identical across panels. Star and triangle markers indicate the best and fastest assemblies, respectively. Dashed lines denote the theoretical optimum.

**Table 2.** Assembly statistics for Ryu, Bcalm2, Hifiasm, and Flye. The misassemblies column shows the number of misassemblies reported by `QUAST`, divided into relocations (flanking sequences align over 1 kbp apart), translocations (flanking sequences align on different chromosomes), and inversions (flanking sequences align on opposite strands).

| Dataset | Tool | Assembly<br>size (Mbp) | N50 (Kbp) | Misassemblies |  |  | Indels | Total<br>contigs |
| --- | --- | --- | --- | --- | --- | --- | --- | --- |
|  |  |  |  | Relocations | Translocation | Inversions |  |  |
| ECOLI | Ryu ★ | 4.69 | 4,640.70 | 1 | 0 | 0 | 5 | 6 |
|  | Ryu ▲ | 4.67 | 4,665.84 | 1 | 0 | 0 | 5 | 4 |
|  | Bcalm2 | 4.53 | 2.19 | 0 | 0 | 0 | 1 | 2,659 |
|  | Hifiasm | 4.67 | 4,665.56 | 1 | 0 | 0 | 4 | 1 |
|  | Flye | 4.64 | 1,182.54 | 1 | 0 | 0 | 3 | 7 |
| YEAST | Ryu ★ | 19.27 | 104.14 | 38 | 26 | 0 | 1,925 | 2,322 |
|  | Ryu ▲ | 19.80 | 65.33 | 47 | 26 | 0 | 1,604 | 1,476 |
|  | Bcalm2 | 10.98 | 1.43 | 0 | 0 | 0 | 355 | 9,753 |
|  | Hifiasm | 33.27 | 138.67 | 463 | 25 | 0 | 4,334 | 890 |
|  | Flye | 11.25 | 759.17 | 0 | 8 | 0 | 238 | 823 |
| HUMAN | Ryu ★▲ | 2,863.07 | 84.33 | 59 | 17 | 4 | 100,672 | 141,087 |
|  | Bcalm2 | 1,249.91 | 1.93 | 0 | 0 | 0 | 157,855 | 827,231 |
|  | Hifiasm | 2,853.07 | 3,967.22 | 177 | 63 | 1 | 157,855 | 1,453 |
|  | Flye | 2,789.13 | 5,953.40 | 108 | 128 | 2 | 190,702 | 2,085 |

**Table 3.** Runtime and peak memory usage for the tools used in the experiments. The instance of `Bcalm2` is the one with the highest N50 value. For `Ryu`, indexing, assembling, and total time are shown for both the best assembly ⋆ and the fastest ▲. The values next to `Bcalm2` correspond to *k* and `-abundance-min`.

| Dataset | Tool |  | Runtime<br>(sec) | Peak<br>memory (GB) |
| --- | --- | --- | --- | --- |
| ECOLI | Ryu | *indexing* | 202 | 0.77 |
|  |  | *assembly* ★ | 261 | 0.10 |
|  |  | *total* ★ | 463 | 0.77 |
|  |  | *assembly* ▲ | 228 | 0.10 |
|  |  | *total* ▲ | 489 | 0.77 |
|  | Bcalm2 | (224, 15) | 97.2 | 6.43 |
|  | Hifiasm |  | 1,799.79 | 16.66 |
|  | Flye |  | 3,279 | 11.43 |
| YEAST | Ryu | *indexing* | 1,555 | 3.11 |
|  |  | *assembly* ★ | 1,428 | 0.44 |
|  |  | *total* ★ | 2,983 | 3.11 |
|  |  | *assembly* ▲ | 1,204 | 0.44 |
|  |  | *total* ▲ | 2,759 | 3.11 |
|  | Bcalm2 | (224, 13) | 378 | 8.11 |
|  | Hifiasm |  | 8,533.65 | 35.63 |
|  | Flye |  | 24,038 | 28.76 |
| HUMAN | Ryu | *indexing* | 7,611 | 13.49 |
|  |  | *assembly* ★▲ | 6,290 | 13.44 |
|  |  | *total* ★▲ | 13,901 | 13.49 |
|  | Bcalm2 | (40,7) | 1,886 | 8.15 |
|  | Hifiasm |  | 3,998.89 | 49.18 |
|  | Flye |  | 74,539 | 104.24 |

### Machine specifications

We performed the experiments in a cluster environment running AlmaLinux 8.10 with kernel Linux 4.18, using a single node per execution: the node uses an Intel(R) Xeon(R) Gold 6248 CPU, with 80 threads and a maximum frequency of 3.9 GHz, with 380GB RAM. All tools were compiled with `GCC 14.2.0` according to the authors’ specifications. We compiled `Ryu` using `-O3`.

## 5.1 Results and discussion

### Effect of *ℓ* and *h* on the assembly

Figure 4 shows how different choices of *ℓ* and *h* affect the results of the assembly. These experiments reveal a tradeoff between contiguity and correctness controlled by the assembly parameters. Additionally, the increasing repeat complexity of the datasets introduces further challenges that influence this tradeoff.

For ECOLI (first row of Figure 4), small values of *ℓ* and *h* produce a larger number of (*ℓ, h*)-tigs, whereas larger values reduce their number. As a consequence, assemblies generated with small *ℓ, h* values are more fragmented and yield smaller N50 values. In contrast, the number of misassemblies follows the opposite trend: smaller values of *ℓ* and *h* produce fewer misassemblies, while larger values increase them.

This behavior can be explained by the way (*ℓ, h*) is used. When *ℓ* and *h* are small, the first node *u* from which the extension and an (*ℓ, h*)-tig can be extended typically has a long label *label*(*u*). This happens because the assembly algorithm requires several extensions before reaching a node *u* such that *f* (*u*) ∈ [*ℓ, h*]. At this stage, no contractions have yet been performed, so the assembly has not introduced misassemblies and the constructed prefix of the (*ℓ, h*)-tig is correct. However, the subsequent chain of contractions and extensions starting from *u* tends to terminates quickly: because the interval [*ℓ, h*] is narrow for small (*ℓ, h*) (by Lemma 1), the frequency of the extending (*ℓ, h*)-tig soon falls outside the allowed range, which limits further extension and results in shorter contigs.

The situation in YEAST is somewhat different (second row of Figure 4). As in ECOLI, increasing (*ℓ, h*) reduces the number of reconstructed sequences. However, the number of misassemblies is minimized for the intermediate choice *ℓ* = 17, *h* = 31. Moving away from this point in either direction increases the number of misassemblies. This more complex behaviour likely reflects the higher structural complexity of the dataset, where unsolved repeats in the reads and homopolymer compression introduce spurious overlaps.

Interestingly, the theoretically optimal choice *ℓ* = 24, *h* = 32, computed using Equation 3, does not minimize the number of misassemblies. However, it does minimize the number of contigs and corresponds to the fastest configuration of `Ryu` on this dataset.

The results for HUMAN (third row of Figure 4) show a pattern similar to ECOLI and YEAST in terms of the number of (*ℓ, h*)-tigs: larger values of *ℓ, h* produce fewer output sequences. However, this metric alone is misleading because the assemblies obtained with larger (*ℓ, h*) also exhibit smaller N50 values, indicating increased fragmentation. The fragmentation when using higher (*ℓ, h*) is due to the lower overage of the HUMAN dataset, 9.57x, and thus we cannot expect the node labels to have higher occurrence frequencies in the reads.

The number of misassemblies in HUMAN shows an opposite pattern compared to ECOLI. Here, larger values of (*ℓ, h*) lead to fewer misassemblies. In this regard, the assembler achieves zero misassemblies for (*ℓ* = 13, *h* = 22), (*ℓ* = 20, *h* = 26), and (*ℓ* = 24, *h* = 32), although these assemblies were highly fragmented.

Overall, these results highlight the trade-off inherent in the choice of (*ℓ, h*). Smaller values favour aggressive extension, producing longer sequences but increasing the risk of misassemblies caused by spurious overlaps. Conversely, larger values restrict extension, which reduces misassemblies but leads to more fragmented assemblies.

### Comparison with other methods

The assembly quality of `Ryu` is comparable to that of long-read assemblers (`Hifiasm`, `Flye`) on simple organisms (ECOLI and YEAST), and substantially better than fixed-order DBG assemblers on complex organisms (HUMAN). At the same time, its computational efficiency remains closer to that of DBG-based tools, highlighting its potential as a lightweight alternative to OLC assemblers. We observe minor differences between the best (⋆) and the fastest (▲) `Ryu` configurations.

Table 2 shows that `Bcalm2` produces a much more fragmented assembly than `Ryu` across all datasets, with very low N50 values and a large number of contigs. By leveraging multiple values of *k*, `Ryu` substantially improves contiguity, producing assemblies with N50 values over 40x higher.

On ECOLI, both `Ryu` and `Hifiasm` reconstruct the genome in a single large contig. `Ryu` produces a few additional short contigs that likely correspond to plasmids or sequencing artifacts, whereas `Flye` splits the genome into several contigs. All methods show similarly low numbers of misassemblies on this dataset.

On the more complex YEAST and HUMAN datasets, `Ryu` produces assemblies that are more contiguous than those of `Bcalm2` but less contiguous than those of `Hifiasm` and `Flye`. However, `Ryu` introduces fewer misassemblies than `Hifiasm` on both datasets and fewer than `Flye` on HUMAN. `Flye` achieves the most contiguous and accurate assembly on YEAST, although this comparison may be biased because the reference genome we used was also assembled using `Flye`.

In terms of computational efficiency (Table 3), `Bcalm2` is the fastest method. Nevertheless, `Ryu` is sub-stantially faster than `Hifiasm` and `Flye` on ECOLI and YEAST, while remaining slower than `Hifiasm` on HUMAN. This comparison is conservative for `Ryu`, since the other assemblers use 24 threads, whereas `Ryu` currently uses four. Moreover, a significant fraction of the runtime in `Ryu` corresponds to index construction.

Finally, `Ryu` consistently uses less memory than full assemblers, illustrating the low memory usage of DBG-based approaches. `Bcalm2` uses even less memory than `Ryu` on the largest dataset because it builds a fixed-order DBG, but this comes at the cost of a substantially increased fragmentation.

### A note on parallel efficiency

When `Ryu` runs in multi-threaded mode with values of *ℓ* and *h* that produce fragmented assemblies, its runtime increases. Parallel execution computes (*ℓ, h*)-tigs across multiple threads while maintaining shared state to avoid generating both DNA strands of the same (*ℓ, h*)-tig independently. In highly fragmented assemblies, threads spend little time extending the contig and more time synchronizing the shared information, leading to high contention. As a result, multi-threaded execution can paradoxically become slower than single-threaded mode. In our experiments, we dealt with this situation by filtering (*ℓ, h*)-tigs shorter than 1000 base pairs: threads discard such contigs and skip the shared-state update. Although this threshold is arbitrary, it had a negligible impact on genome coverage (see Table 2).

## 6 Conclusions and further work

We presented a theoretical framework for assembling long reads using voDBGs. Our experiments show that `Ryu` achieves a strong compromise between contiguity, correctness, and efficiency, significantly improving over fixed-order DBGs while remaining lighter than long-read assemblers. These results suggest that voDBG-based assemblers can offer a practical alternative to expensive OLC methods.

Further improvements could make our approach a practical full-fledged *de novo* assembler. Advances in read indexing and misassembly detection may improve both efficiency and accuracy, while ongoing progress in algorithms for constructing and indexing large BWTs is likely to enable more scalable implementations. Additionally, dynamically adjusting the interval [*ℓ, h*] based on local read features could help reduce misassemblies, although identifying the most informative signals remains a challenge. On the other hand, scaffolding strategies that exploit the information in the read index may help reduce fragmentation. Finally, extending the framework to polyploid genomes could enable efficient reconstruction of more complex genomes.

## Acknowledgments

We used AI tools to improve clarity and tighten the text to fit the page limit.

## A Appendix

**Table A1.** Comparison between all the instances of Ryu with Hifiasm, Flye and the best two instances of Bcalm2. In bold, the theoretical optimal values of (*l, h*) for the given genome. With ⋆ (resp. ▲) we refer to the best (resp. fastest) assembly.

| Dataset | Tool | Parameter | N50<br>(Kbp) | Misassemblies | Largest<br>Contig<br>(Kbp) | Total<br>Contigs |
| --- | --- | --- | --- | --- | --- | --- |
| ECOLI | Bcalm2 | 128,13 | 1.20 | 0 | 7.07 | 3,301 |
|  |  | 224,15 | 2.19 | 0 | 15.12 | 2,659 |
|  | Flye |  | 1,182.54 | 1 | 2,087.87 | 7 |
|  | Hifiasm |  | 4,665.56 | 1 | 4,665.56 | 1 |
|  | Ryu | 9,16 | 13.29 | 0 | 25.67 | 11 |
|  |  | 9,17 | 13.29 | 0 | 25.67 | 10 |
|  |  | 13,22 | 4,654.42 | 1 | 4,654.42 | 8 |
|  |  | 13,25 | 24.55 | 0 | 24.55 | 7 |
|  |  | 17,28 | 4,640.70 | 1 | 4,640.70 | 6 |
|  |  | 17,31 ★ | 4,665.84 | 1 | 4,665.84 | 4 |
|  |  | **18,25 ▲** | 4,640.70 | 1 | 4,640.70 | 6 |
|  |  | 20,26 | 4,640.70 | 1 | 4,640.70 | 6 |
|  |  | 24,32 | 4,640.70 | 1 | 4,640.70 | 5 |
| YEAST | Bcalm2 | 224,13 | 1.43 | 0 | 9.66 | 9,753 |
|  |  | 256,15 | 0.89 | 1 | 15.85 | 22,545 |
|  | Flye |  | 759.17 | 8 | 1,471.70 | 23 |
|  | Hifiasm |  | 138.67 | 488 | 1,357.40 | 890 |
|  | Ryu | 9,16 | 47.43 | 87 | 505.03 | 3,790 |
|  |  | 9,17 | 58.66 | 85 | 590.06 | 3,673 |
|  |  | 13,22 | 70.44 | 70 | 391.44 | 2,508 |
|  |  | 13,25 ★ | 104.14 | 64 | 626.48 | 2,322 |
|  |  | 17,28 | 71.48 | 66 | 381.41 | 1,825 |
|  |  | 17,31 | 100.67 | 60 | 511.24 | 1,675 |
|  |  | 20,26 | 60.30 | 77 | 322.16 | 1,848 |
|  |  | 24,32 | 64.83 | 74 | 322.15 | 1,484 |
|  |  | **23,33 ▲** | 65.33 | 73 | 322.15 | 1,476 |
| HUMAN | Bcalm2 | 32,7 | 1.73 | 0 | 25.15 | 827,231 |
|  |  | 40,7 | 1.93 | 0 | 28.41 | 846,616 |
|  | Flye |  | 5,953.40 | 238 | 26,830.28 | 2,085 |
|  | Hifiasm |  | 3,967.22 | 241 | 33,485.86 | 1,453 |
|  | Ryu | 5,9 ★▲ | 84.33 | 80 | 674.80 | 141,087 |
|  |  | 9,16 | 18.72 | 11 | 169.04 | 221,144 |
|  |  | 9,17 | 19.03 | 74 | 169.04 | 212,741 |
|  |  | 13,22 | 6.04 | 0 | 75.77 | 103,080 |
|  |  | 13,25 | 6.46 | 33 | 78.88 | 88,931 |
|  |  | **15,20** | 3.92 | 0 | 53.08 | 61,922 |
|  |  | 17,28 | 3.12 | 2 | 53.08 | 34,444 |
|  |  | 17,31 | 3.32 | 5 | 62.87 | 31,029 |
|  |  | 20,26 | 2.53 | 0 | 47.72 | 25,504 |
|  |  | 24,32 | 2.22 | 0 | 35.24 | 20,426 |

## Footnotes

1 https://github.com/ddiazdom/Ryu, commit 0eb064a

2 https://github.com/GATB/bcalm, commit fe8a8c5

3 https://github.com/mikolmogorov/Flye, commit 886b8c1

4 https://github.com/chhylp123/hifiasm, commit ec9a8b2

